# GENE EXPRESSION WITHIN A HUMAN CHOROIDAL NEOVASCULAR MEMBRANE USING SPATIAL TRANSCRIPTOMICS

**DOI:** 10.1101/2023.06.16.544770

**Authors:** Andrew P. Voigt, Nathaniel K. Mullin, Emma M. Navratil, Miles J. Flamme-Wiese, Li-Chun Lin, Todd E. Scheetz, Ian C. Han, Edwin M. Stone, Budd A. Tucker, Robert F. Mullins

**Author notes:** Correspondence: Robert F. Mullins 4130 MERF 375 Newton Rd Iowa City, IA 52242 USA. **FUNDING:** National Eye Institute (R01EY033308, R01EY033331, P30EY025580, F30EY034009, F30EY031923); National Institute of General Medical Sciences (T32GM139776); The Edward N. & Della L. Thome Memorial Foundation; The Elmer and Sylvia Sramek Charitable Trust. **CONFLICT OF INTEREST:** No authors declare competing interests. **AUTHOR CONTRIBUTIONS:** APV, TES, ICH, EMS, BAT, and RFM performed study concept and design; APV, NKM, MJWF performed development of methodology; APV and RFM performed writing, review and revision of the paper; APV, NKM, EMN, MJFW, LL provided acquisition, analysis and interpretation of data, and statistical analysis; TES, ICH, EMS, BAT, and RFM provided technical and material support. All authors read and approved the final paper. **DATA AVAILABILITY:** All raw sequencing and imaging data have been deposited in the Gene Expression Omnibus database GSE234047.

## Abstract

Macular neovascularization is a relatively common and potentially visually devastating complication of age-related macular degeneration. In macular neovascularization, pathologic angiogenesis can originate from either the choroid or the retina, but we have limited understanding of how different cell types become dysregulated in this dynamic process. In this study, we performed spatial RNA sequencing on a human donor eye with macular neovascularization as well as a healthy control donor. We identified genes enriched within the area of macular neovascularization and used deconvolution algorithms to predict the originating cell type of these dysregulated genes. Within the area of neovascularization, endothelial cells were predicted to increase expression of genes related to Rho family GTPase signaling and integrin signaling. Likewise, VEGF and TGFB1 were identified as potential upstream regulators that could drive the observed gene expression changes produced by endothelial and retinal pigment epithelium cells in the macular neovascularization donor. These spatial gene expression profiles were compared to previous single-cell gene expression experiments in human age-related macular degeneration as well as a model of laser-induced neovascularization in mice. As a secondary aim, we also investigated spatial gene expression patterns within the macular neural retina and between the macular and peripheral choroid. We recapitulated previously described regional-specific gene expression patterns across both tissues. Overall, this study spatially analyzes gene expression across the retina, retinal pigment epithelium, and choroid in health and describes a set of candidate molecules that become dysregulated in macular neovascularization.

## INTRODUCTION

Age-related macular degeneration (AMD) is a common cause of blindness in which the central retina (termed the macula) degenerates, leading to a loss of high acuity center vision. Advanced stages of AMD include a progressive deterioration (geographic atrophy) of the retinal pigment epithelium and choriocapillaris vascular layer and/or abnormal new blood vessel growth called macular neovascularization (MNV). In MNV, pathologic angiogenesis can result in severe and often rapid vision loss due to retinal damage from hemorrhage and vascular leakage. Thanks in large part to developments in imaging, MNV can be reproducibly categorized as Type I (formerly “occult” choroidal neovascularization) that arises from the choroidal vasculature and lies between the retinal pigment epithelium (RPE) and Bruch’s membrane, Type II (formerly “classic” choroidal neovascularization) that similarly arises from the choroid but that grows into the subretinal space, and Type III MNV that originates from the retinal vasculature rather than the choroid (1). MNV can be exudative, with serous fluid seeping past the compromised RPE and damaging the overlying photoreceptors. By contrast, non-exudative MNV has been conjectured to be protective against geographic atrophy by re-establishing a choriocapillaris-like layer to meet the metabolic demands of the retina (2).

The application of anti-VEGF agents has been remarkably effective at treating MNV, and millions of patients have had tremendous positive visual outcomes with this therapy (3). In most patients, anti-VEGF is effective at resolving subretinal and intraretinal fluid (4). However, in some patients, the response to anti-VEGF drugs may be limited or wear off over time (5-8), and the exact mechanism(s) of this attenuated response remain poorly understood (9). The visual consequences of MNV and the frequency of associated treatment (often monthly clinic-based injections) significantly impacts affected individuals’ quality of life (10). An improved understanding of the pathophysiologic mechanisms involved in MNV may inform the development of targeted interventions for those with limited response to anti-VEGF or those that develop geographic atrophy despite treatment (11).

Human neovascular membranes removed during surgery (12) and endothelial cells from the mouse model of laser induced choroidal neovascularization (13) have been previously studied at the transcriptional level using bulk and single-cell RNA sequencing, respectively. We recently evaluated choroidal gene expression at the single-cell level in a series of eyes from donors with atrophic AMD, age-matched controls, and two eyes with MNV (14). Such approaches highlight dysregulated gene expression in MNV. However, a limitation of transcriptomic studies to date is that the spatial location of each cell is not captured, even when performed at the single cell level. For example, when tissue from an MNV is dissociated, it is not possible to discriminate between endothelial cells within the neovascular membrane versus those in the adjacent and underlying choroid. To detect transcriptomic features within a MNV with greater specificity, spatial analysis of gene expression patterns offers greater potential.

In the current study, we performed spatial RNA sequencing on two human donor eyes, one from an unaffected control and one eye from a patient with Type I MNV. Consistent with our expectations, gene expression patterns in the normal retinal outer nuclear, inner nuclear, and ganglion cell layers were highly enriched for photoreceptor, Müller cell, and ganglion cell transcriptomic profiles, respectively. We further identified gene signatures spatially enriched in the neovascular membrane compared to normal choroid. Such spatial profiles were compared to previous single-cell gene expression patterns in human AMD endothelial cells (14) and models of laser-induced neovascular endothelial cells in mice (13). As a secondary aim, we identified gene expression patterns that varied spatially within the macular neural retina and compared these results to previous regional single-cell RNA sequencing studies (15).

## METHODS

### Human donor eyes

All donor tissue used in this study was recovered by the Iowa Lions Eye Bank with full consent from the next of kin for use in research and evaluation of donor clinical records. All studies were performed in compliance with the Declaration of Helsinki. For the spatial transcriptomics experiments, eyes from two donors were employed. The first was from a 78-year-old Caucasian female who was pseudophakic in both eyes but with an otherwise unremarkable ophthalmic history. The second was from an 85-year-old Caucasian female with neovascular AMD in both eyes. From the right eyes of both donors, a 4 mm diameter macular punch was collected centered on the fovea centralis, as was a 4 mm peripheral punch approximately 12 mm superotemporal from the macula. Both central and peripheral punches (spanning retina, choroid and sclera) were placed in optimal cutting temperature compound (OCT; Sakura) and were rapidly snap frozen a in liquid nitrogen bath before storage at - 80°C.

### Spatial RNA library generation

Cryostat sections were collected at a thickness of 10µm onto a precooled Visium slide (10X Genomics, Pleasanton CA). Visium slides with the sections were fixed, stained, and imaged with hematoxylin and eosin using a 20X objective on an Olympus BX61 Upright Microscope. Tissue was then permeabilized for 12 minutes, which was established as an optimal permeabilization time based on tissue optimization time-course experiments. The poly-A mRNAs from the sections were released and captured by the poly(dT) primers precoated on the slide, including spatial barcodes and unique molecular identifiers (UMIs). After reverse transcription and second strand synthesis, the amplificated cDNA samples from the Visium slides were transferred, purified, and quantified for library preparation.

Reverse transcription, second strand synthesis and denaturation, and cDNA amplification were performed as described by the manufacturer (10X Genomics Spatial Gene Expression Protocol CG000239 Revision F). The final libraries were pooled and sequenced on the NovaSeq 6000 platform (Illumina, San Diego, CA) generating 100-bp, paired end reads.

### Bioinformatic analysis

FASTQ files were generated from base calls with the bcl2fastq software (Illumina, San Diego CA). FASTQ files were mapped to the human GRCh38 reference genome with spaceranger (v1.3.1). Raw FASTQ files and processed data matrices have been deposited in the Gene Expression Omnibus database (GSE234047). Count data from each slide was loaded into Seurat (16) with the Read10X_Image() function. All sections underwent normalization with the SCTransform() function. Data from all sections were merged and subsequently integrated into one dataset before PCA, clustering, and uniform manifold approximation and projection (UMAP) dimensionality reduction.

In addition, we performed a deconvolution analysis with CSIDE (17) to estimate the proportion of different cell types contributing RNA to each spot using our previous single-cell expression studies in the human retina (15) and RPE-choroid (18) as references for cell type label transfer. Using the deconvoluted weights, differential expression analysis was completed with the run.CSIDE() function in the spacexr R package (v2.0.1). To increase the number of candidates genes available for comparison with other datasets, CSIDE was run in the full (non-doublet) mode, the FDR was set to 0.9, and the cell type threshold was set to 10. Three sets of differential analyses were completed. First, gene expression was compared between spots overlapping the MNV to spots outside of the MNV or in control tissues. Covariates for the analysis included cell type deconvolution and the percentage of spot area overlapping with the MNV (see below). MNV-enriched genes were compared to previous single-cell RNA sequencing studies of laser-induced choroidal neovascularization in mice (GSE77199, (13)) and two human donors with MNV (GSE183320, (14)). Likewise, endothelial and RPE genes with a log_2_FC enrichment > 0.5 in the MNV sample were used as input for ingenuity pathway analysis. Second, gene expression was spatially evaluated along a gradient of distance to the fovea. We measured the distance between each spot and the center of the section and scaled this distance from 0 to 1. This distance and cell type deconvolution were used as the covariates for differential expression analysis, and results were compared to a single-cell RNA sequencing study of foveal (1 mm) versus parafoveal (4 mm) human retina (GSE169046, (15)). Third, we compared gene expression between macular and peripheral choroidal deconvoluted spots. Spots were manually categorized as macular or peripheral prior to differential expression.

### Manual Classification of Spots

The LoupeBrowser (10X Genomics, Pleasanton CA) was used to manually annotate spots originating in macular versus peripheral sections and overlying retinal versus RPE/choroid tissue. These labels were added to the meta data of each section within Seurat. In addition, the MNV was manually traced and intersected with overlying spots, and ImageJ was used to quantify the proportion of each spot that overlapped the neovascular membrane. Spots overlying the sclera were excluded from all analyses.

## RESULTS

### Clinical and Histologic Evaluation of Eyes Used in Transcriptomic Experiments

Ocular tissue from two human donors was acquired for spatial transcriptomic experiments. The first donor was a 78-year-old Caucasian female with a history of cataract removal but no other ocular history. The posterior pole of this control donor appeared grossly normal with normal foveal and choroidal pigmentation (**Figure 1A**). The second donor was an 85-year-old Caucasian female with a >5-year year history of MNV in both eyes. Visual acuity was 20/20 in the right eye and 20/250 in the left eye, with records of an involuted choroidal neovascular membrane in both eyes for the last 5 years. A yellow elevation of the retina with areas of both hypopigmentation and hyperpigmentation was observed by gross examination in both maculas of the MNV donor (**Figure 1B**, right eye shown). Infrared reflectance imaging and optical coherence tomography acquired one month prior to death revealed the presence of superior reticular pseudodrusen and a very broad, low-lying fibrovascular pigment epithelial detachment with “double-layer sign” (19) within the macula, consistent with type 1 MNV (**Figure 1C-D**). Hematoxylin and eosin staining of the control (**Figure 1E**) macula demonstrated normal photoreceptor outer segments, RPE, and choriocapillaris. In contrast, the MNV donor had a prominent type I neovascular membrane between Bruch’s membrane and the retinal pigment epithelium (**Figure 1F**).

**Figure 1:**
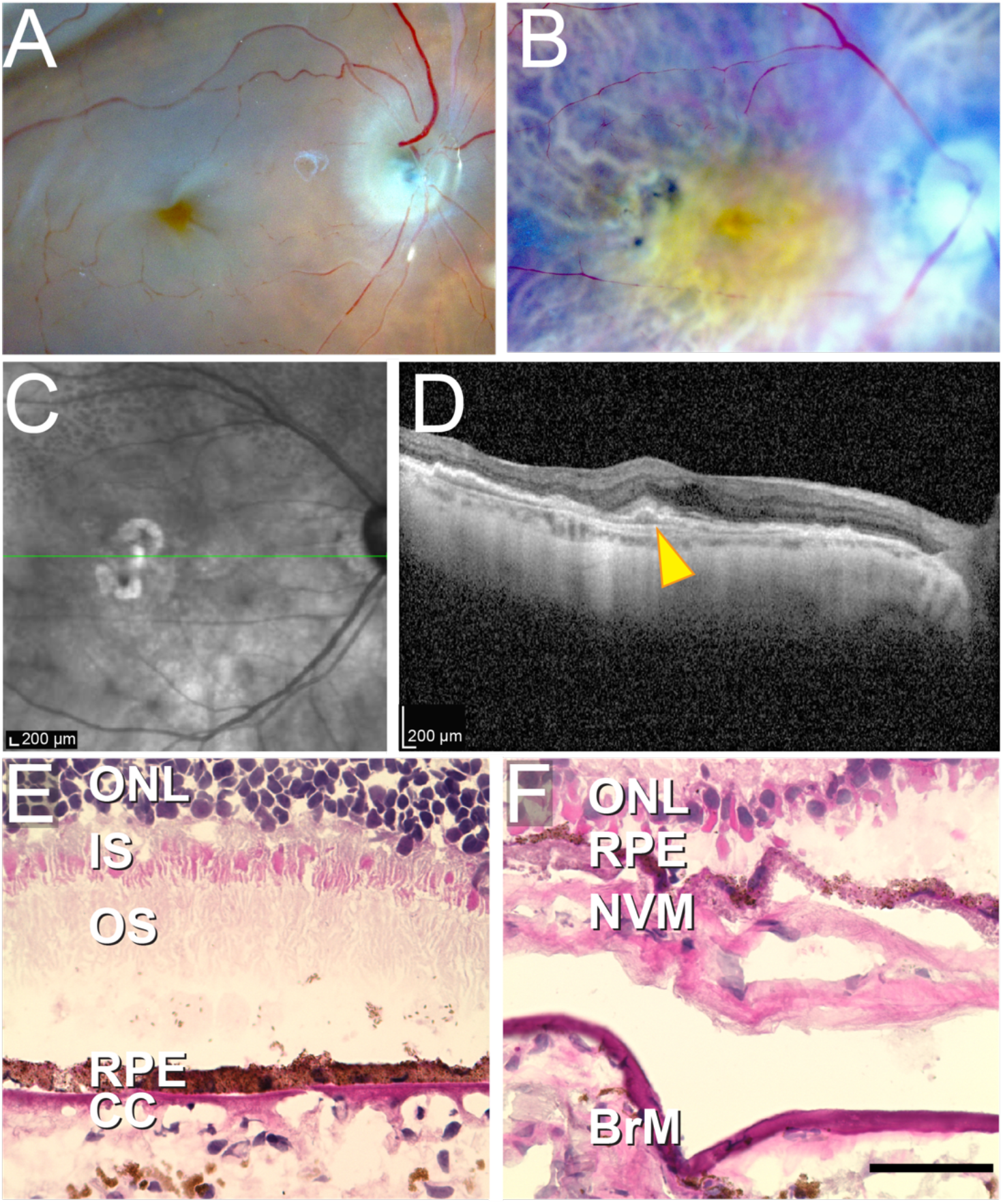
Gross, clinical and histological characterization of eyes used for spatial transcriptomic experiments. Gross photos from the control (**A)** and MNV (**B**) donor after dissection. **A.** 4-mm punch was acquired centered on the fovea and approximately 12 mm superotemporal to the macula. While the macula of the healthy control donor appeared grossly normal (**A**), the MNV donor macula (**B**) demonstrated a disc-diameter area of elevated yellow material and atrophy. Near infrared scan (**C**) and optimal coherence tomography scan (**D**) of the right eye from the patient with macular neovascularization showing a broad, low-lying fibrovascular pigment epithelial detachment with “double-layer sign” (arrowhead). Reticular pseudodrusen (dark rounded profiles on near infrared scan) were noted superior to the lesion along the arcade vessels. Histological macular tissue sections from the control (**E**) and MNV (**F**) donor stained with hematoxylin-eosin. Note the thinning of the outer nuclear layer and fibrovascular membrane in the MNV eye. ONL = outer nuclear layer; IS, inner segments; OS, outer segments; NVM, neovascular membrane; BrM, Bruch’s membrane, CHO = choroid.

### Spatial RNA Sequencing Overview

A total of five tissue sections (n = 3 MNV, n = 2 control) were acquired for spatial RNA sequencing (**Figure 2A**). Macular and peripheral tissues were embedded in the same OCT block and sectioned together onto each capture area, and each section consisted of macular retina, macular RPE/choroid, and peripheral RPE/choroid. The tissue sections were fixed in methanol, stained with hematoxylin and eosin, and imaged (**Figure 2B, SI Figure 1**). Each slide was subsequently permeabilized and cDNA libraries were prepared and sequenced. After mapping to the human genome, unique barcodes were used to spatially link each read to the x/y coordinate of a spot on the gene expression slide.

**Figure 2:**
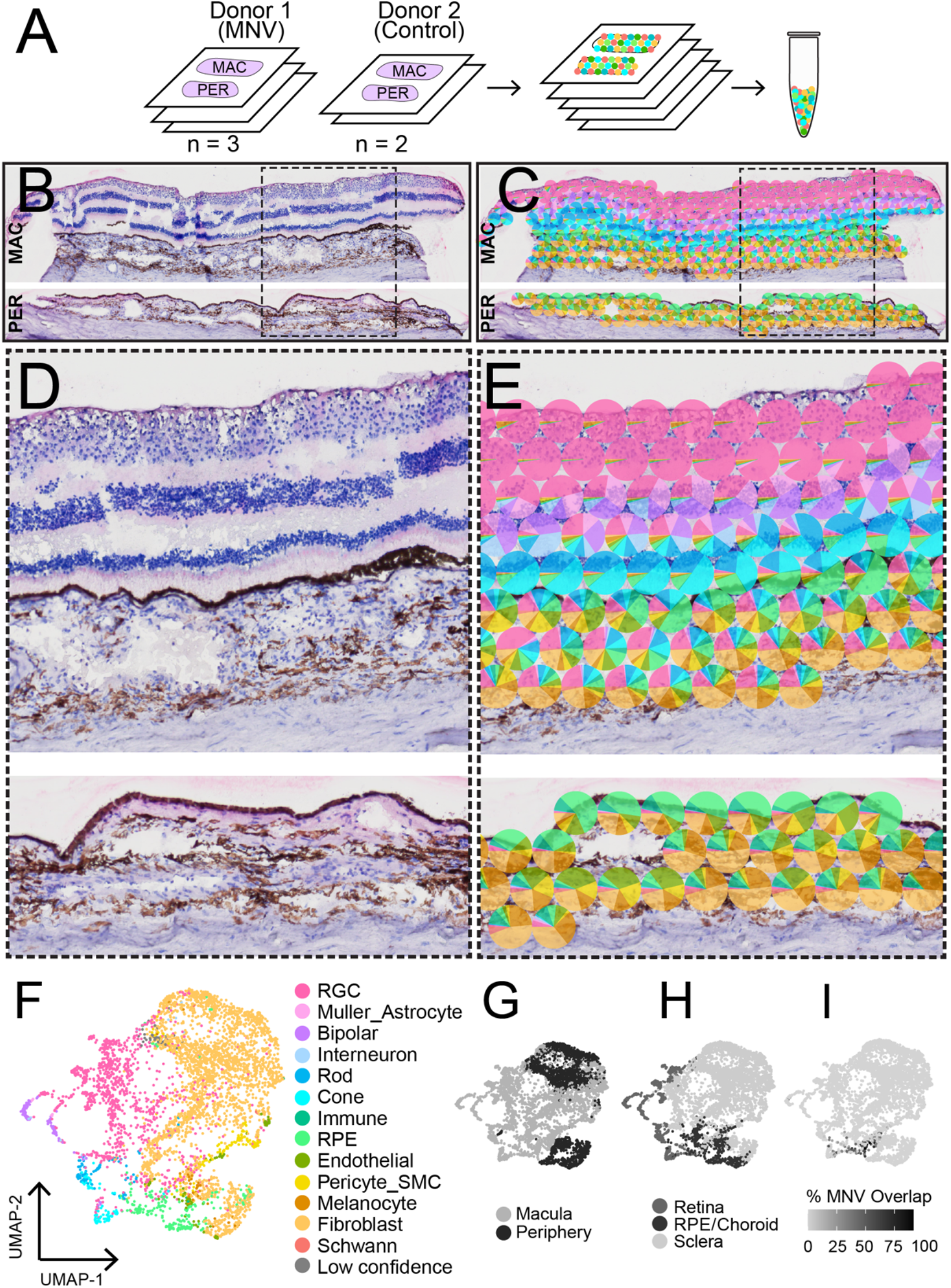
Spatial RNA sequencing of a control and MNV human donor macula. **A.** Experimental overview: tissue sections from a control donor (n = 2 sections) and an MNV donor (n = 3 sections) were prepared for spatial RNA sequencing. After fixation and imaging, sections were permeabilized and cDNA libraries were prepared. Hematoxylin and eosin staining of one section of the control donor (**B**). The RNA contribution from retinal, RPE, and choroidal cell types was estimated in each spatially barcoded spot (**C**) and displayed as a pie-chart. (**D**) and (**E**) show a magnified view of the spot area in the macular (top) and peripheral (bottom) sections. Spots overlying the sclera were excluded from this visualization. **F.** Uniform manifold approximation and projection (UMAP) dimensionality reduction was applied to visualize clusters of spots with similar gene expression profiles. Each point represents the multidimensional transcriptomic profile from one spot, and points are colored according to the maximally predicted RNA contribution in each spatially barcoded spot. Clustering was visualized according to region (**G**), tissue (**H**), and percentage of the spot area that overlaps with the MNV membrane (**I**).

Consistent with the cell types known to reside in each retinal layer, the different retinal layers demonstrated distinct gene expression patterns. For example, spots in the ganglion cell layer contained high amounts of *NEFL*, spots overlying the outer nuclear layer were enriched in the cone-specific phosphodiesterase *PDE6H*, and spots overlying the RPE were enriched in *LRAT* (**SI Figure 2**). The diameter of each barcoded spot was 55 µm, which typically covered multiple different cells within the densely cellular retina, RPE, and choroid. To overcome this technical limitation, we next estimated the RNA contribution of different cell types within each spot using a deconvolution algorithm and previously published retinal (15) and choroidal (14) single-cell RNA sequencing studies as reference datasets (**Figure 2C**). Indeed, predicted cellular compositions were unique between the different retinal layers (**Figure 2D-E**), and distinct tiers of spots highly enriched for retinal ganglion cells, bipolar cells, photoreceptors, and RPE cells were observed. Spots with similar multidimensional transcriptomic profiles were clustered and classified based on which cell type was predicted to contribute the most RNA to each spot (**Figure 2F**). Clustering was visualized according to region, tissue, and percentage of the spot area that overlapped with the MNV membrane (**Figure 2G - I**). As the spots have a relatively large diameter, it is unsurprising that spots often clustered by region (macula versus peripheral) (**Figure 2G**) and/or tissue (retina, RPE, and choroid) (**Figure 2H**).

### Gene Expression Changes in the MNV Lesion

As the primary aim of the study, we investigated gene expression differences between spots within the MNV lesion versus non-MNV RPE and choroid. First, we manually traced the MNV membrane to visualize how the spots overlapped with the pathology (**Figure 3 A-C**). A total of 144 barcoded spots partially overlapped with the lesion across the three sections. However, due to the relatively large spot diameter of 55 microns, no spot was entirely contained in the MNV lesion. We used ImageJ to calculate the percentage of spot area overlapping the traced membrane (**Figure 3D**), which ranged from <1% to 91%. Next, we assessed the deconvoluted cell type contributions of spots overlapping the membrane (**Figure 3E**). RPE cells were predicted to contribute the most RNA to spots overlapping the MNV, followed by retinal ganglion cells, cone photoreceptor cells, endothelial cells, and pericyte/smooth muscle cells. The presence of inner retinal transcripts within the MNV-overlapping spots may represent non-specific diffusion (20) (which can be due to over permeabilization (21)) of RNA beyond the cell margins or imprecise deconvolution predictions.

**Figure 3:**
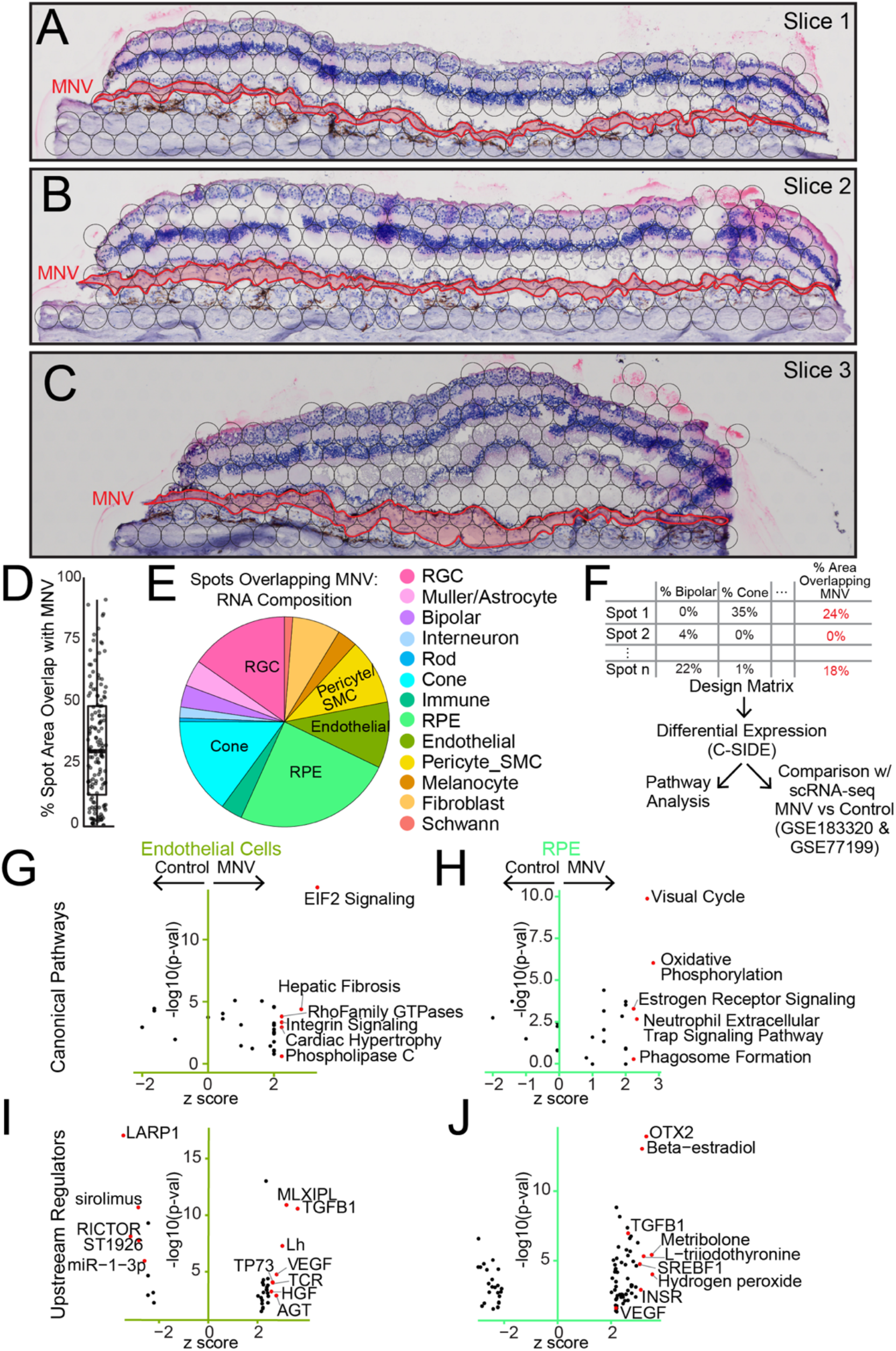
Gene expression in a human macular neovascular membrane. **A-C**. The three macular sections from the MNV donor. The neovascular membrane (red outline) is traced in each tissue section and overlapping spot areas (black circles) are visualized. Note that the neovascular membrane spanned the entire section in each case. **D**. The percent area that each spot overlapped the MNV membrane was calculated. A total of 144 spots overlapped the lesion, ranging from <1% to 91% overlap. **E**. For all spots overlapping the MNV, the relative cell type composition was calculated. **F.** Design of differential expression analysis. Cell type deconvolution and percent area overlap with the MNV were used as covariates for differential expression analysis. **G-J**. After identifying genes enriched in MNV spots, ingenuity pathway analysis was used to identify canonical pathways (**G, H**) and upstream regulators (**I, J**) of genes predicted to originate from endothelial cells and RPE cells.

Each spot overlying the MNV lesion was composed of multiple underlying cell types and had varying overlap with the neovascular membrane. To account for the multiple sources of variation, we used the CSIDE parametric differential expression algorithm (17) to identify MNV-enriched genes. CSIDE uses cell type mixtures (identified by deconvolution) as covariates to detect genes whose expression is related to the continuous explanatory variable of spot-MNV overlap area (**Figure 3F**). CSIDE then provides differential expression results split for each individual cell type (**SI File 1**).

Gene expression changes in neovascular endothelial cells and the overlying RPE were of particular interest, and we set out to understand the functional significance of MNV gene expression dysregulation. As such, we performed pathway enrichment analysis to find canonical pathway terms (**Figure 3G-H**) of genes enriched in MNV-overlapping spots predicted to originate from endothelial cells and RPE cells (**SI File 1**). Endothelial genes from spots overlapping the MNV lesion were enriched in pathways terms related to Rho Family GTPase signaling and Integrin signaling (**Figure 3G**). Rho GTPases can regulate angiogenesis by modulating *VEGF* expression levels (22). The genes *CFL1* and *ITGB1*, which encode the actin binding protein cofilin and the beta-1 integrin subunit, were increased in neovascular endothelial cells. These gene products are involved in Rho signal transduction and have been implicated in endothelial invasion and angiogenesis (23, 24). Similarly, integrins can stimulate pathological angiogenesis by transducing signaling through VEGF and other growth factors (25). The RAS family genes *RHOC* (which encodes a protein which colocalizes with integrin α5β1) and *RAP1B* (which encodes a product that colocalizes with integrin αvβ₃) have been shown to regulate VEGF activation (22, 26), and both genes were increased in MNV endothelial cells. RPE genes from spots overlapping the membrane were enriched in pathway terms related to ERK/MAPK signaling (**Figure 3H**), which promotes endothelial cell sprouting and angiogenesis (27). For example, MNV-overlapping spots demonstrated increased expression of *ITGAV* and *ITGB8* (two genes highly expressed by the RPE), which encode integrin components that regulate ERK/MAPK signaling and have been associated with angiogenesis and blood vessel network formation (28, 29).

Similarly, we searched for predicted upstream regulatory factors that could drive the observed gene expression differences in MNV-overlapping spots (**Figure 3 I-J**). Ingenuity analysis identified VEGF and TGFB1 pathways as upstream regulators of MNV gene expression changes predicted to originate from endothelial cells and RPE cells. VEGFs are the central regulators of angiogenesis and are strongly implicated in MNV (30), and the gold standard for treating neovascular AMD is anti-VEGF drugs (e.g., (31)). Likewise, TGFB signaling has complex roles in modulating angiogenesis and vessel formation (32) and has been previously shown to participate in choroidal neovascularization in mice (33). Overall, these data suggest that the observed MNV-associated endothelial and RPE cell gene expression patterns may be driven by known growth factors central to MNV.

### Comparing Neovascular Membrane Enriched Genes with Single-Cell Studies

To date, there have been two studies that explored gene expression changes in neovascular choroidal endothelial cells at the single-cell level. First, Rohlenova et al used a laser injury model to induce choroidal neovascularization in mice before single-cell RNA sequencing of enriched endothelial cells, using the contralateral eye of each mouse as a control (13). Second, our group performed single-cell RNA sequencing on 21 human donor choroids, which included 10 control donors and 2 donors with MNV (14). In both studies, even the pathologic samples contained non-neovascular endothelial cells alongside neovascular endothelial cells. However, the lack of spatial barcoding in these studies makes it difficult to positively identify the endothelial cells within the neovascular membrane and identify unique gene expression features specific to these cells. Despite these limitations, we hypothesized that many of the endothelial MNV-enriched genes identified by the current spatial study would also be enriched in the neovascular samples from the previous single-cell investigations despite the known mixed population of normal and pathological endothelial cells.

To test this hypothesis, we identified the top ten MNV and control enriched genes predicted to originate from endothelial cells in the current study and compared these results to reprocessed single-cell investigations (**Figure 4**). Overall, 70% of single-cell RNA sequencing log_2_ fold-changes were in the same direction as the current spatial study. In the current study, the endothelial gene with the highest MNV log_2_ fold-change was *IL6*. IL6 is a potent stimulator of vessel sprouting and angiogenesis (34), and *IL6* RNA levels have been previously shown to increase in choroidal macrophages in a laser model of choroidal neovascularization in mice (35). Previous single-cell RNA sequencing data (14) suggests *IL6* is expressed at the RNA level by choroidal endothelial cells, inflammatory macrophages, fibroblasts, and pericytes. Unfortunately, due to the large spot diameter in the current study, it is challenging to distinguish if endothelial cells, the closely associated inflammatory macrophages, or both cell types are responsible for the increased *IL6* within the MNV of this donor.

**Figure 4:**
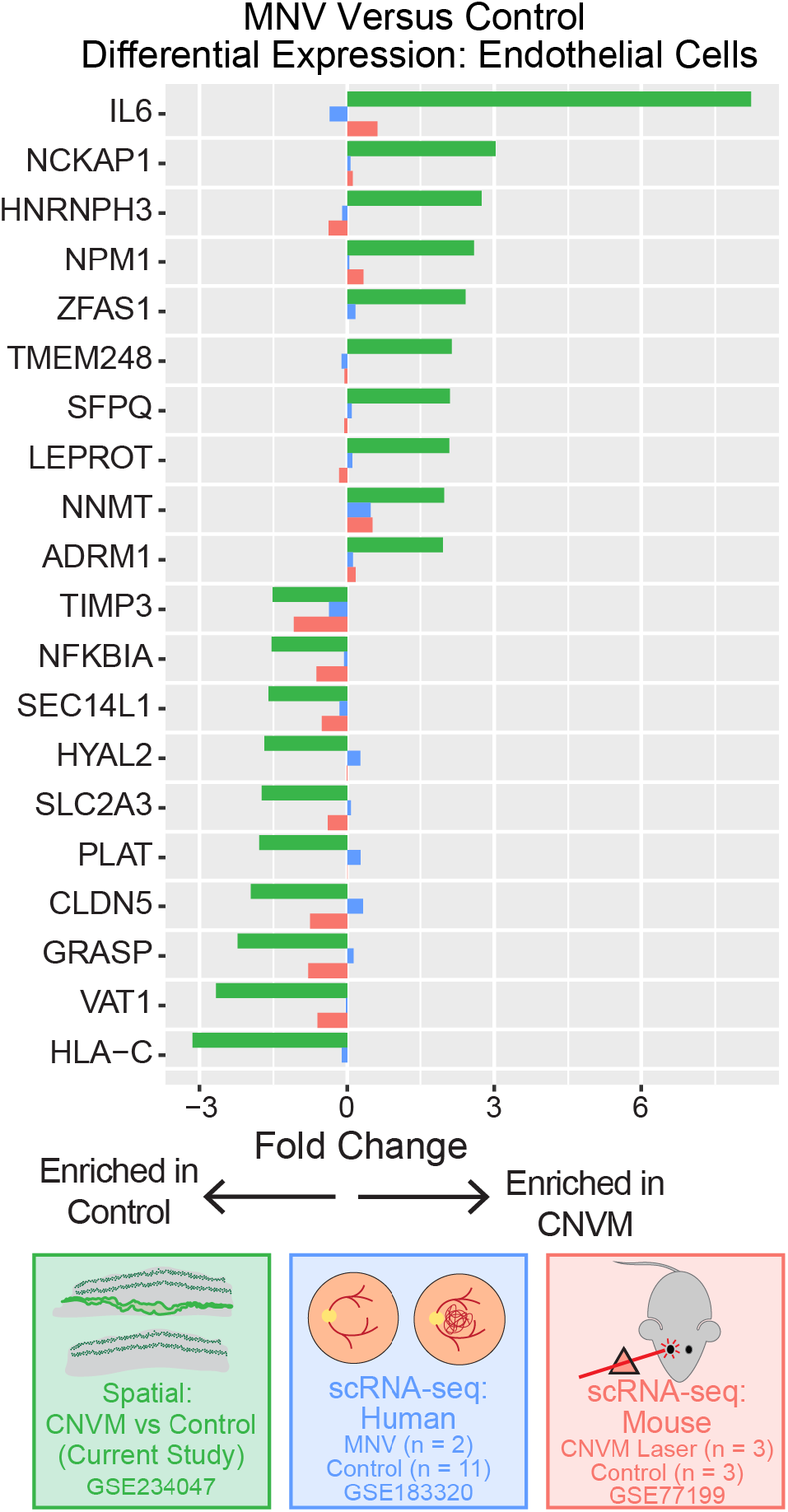
Comparison of MNV-enriched genes in endothelial cells with previous single-cell RNA sequencing studies. Genes upregulated and downregulated in MNV-associated spots according to spatial RNA sequencing (green, current study) were compared to a previous single-cell RNA sequencing study of two human donors with MNV (blue) and a mouse model of choroidal neovascularization (red). Positive log_2_ fold-changes are associated with increased expression in the neovascular endothelial cells.

### Regional Gene Expression in the Neural Retina and RPE/Choroid

We next used spatial RNA sequencing to investigate regional patterns of gene expression within neural retina, RPE, and choroid. Our spatial sections captured macular (but not peripheral) neural retina, so we first investigated how gene expression varies within the macula. Each 4 mm punch used for this investigation was centered on the fovea. While it is unlikely that the fovea was perfectly centered, we assumed that the center of the section would be closer to the fovea and that the edges of the section would be further from the fovea. Therefore, we measured the distance of each spot to the center of each section, and we scaled this distance from 0 (close to the fovea) to 1 (far from the fovea). Next, we performed differential expression with CSIDE to identify genes with varied regional expression patterns. Like the MNV differential expression, CSIDE used cell type mixtures (identified by deconvolution) as covariates to detect genes whose expression is related to the continuous explanatory variable of distance from the section center point (**SI File 2**).

We then compared these differential expression results of cone photoreceptor and Müller cells to a previous single-cell investigation of foveal (1 mm) versus parafoveal (4 mm) neural retina from four human donors (15). We hypothesized that genes previously identified to vary regionally in the neural retina would demonstrate similar enrichments in the current spatial dataset (**Figure 5**). Overall, 83% of cone photoreceptor and 93% of Müller cell log_2_ fold-changes were in the same direction as the previous single-cell RNA sequencing study.

**Figure 5:**
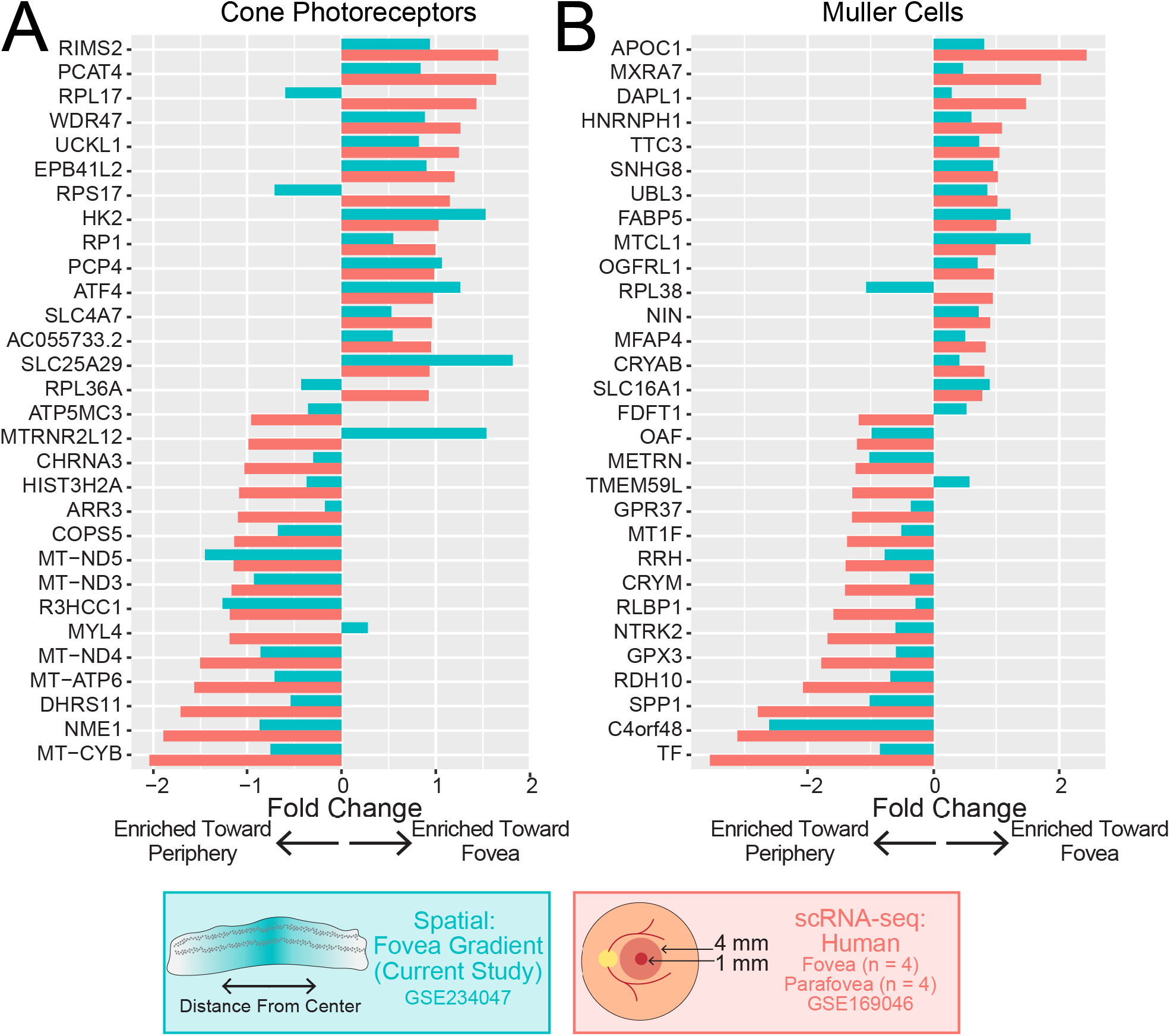
Comparison of regional retinal gene expression differences between spatial and single-cell RNA sequencing studies. Differential expression was completed to compare genes enriched toward the center and towards the periphery of each section. For macular sections, the distance between each spot and the center of the slide was calculated and scaled from 0 to 1. Cell type deconvolution and distance from the center were used as covariates for differential expression analysis. For cone photoreceptor cells (**A**) and Muller cells (**B**), the top differentially expressed genes between the fovea and parafovea were identified from a previous single-cell RNA sequencing study. Differential expression results were compared between this previous study (red) and the current spatial study (blue). Positive log_2_ fold-changes are associated with increased expression towards the center of the retina.

In addition to the neural retina, each tissue section contained both macular and peripheral RPE/choroid. We next compared how gene expression differed between the macula and periphery of the control donor. Similar to the neural retina investigation, we performed differential expression with CSIDE using cell type mixtures (identified by deconvolution) as covariates to detect genes whose expression is related to the binary explanatory variable macular or peripheral origin (**SI File 3**). We then compared these spatial enrichments of RPE and endothelial cells to a previous single-cell RNA sequencing study of macular versus peripheral human RPE/choroid (36). We hypothesized that genes previously identified to vary between these two regions in the RPE/choroid would demonstrate similar enrichments in the current spatial dataset (**Figure 6**). A total of 75% of RPE cell and 75% of endothelial cell log_2_ fold-changes were in the same direction as this previous single-cell RNA sequencing study. Overall, these data suggest that even with large spot diameters, spatial RNA sequencing can detect regional-specific gene expression patterns within the heterogenous neural retina and RPE/choroid.

**Figure 6:**
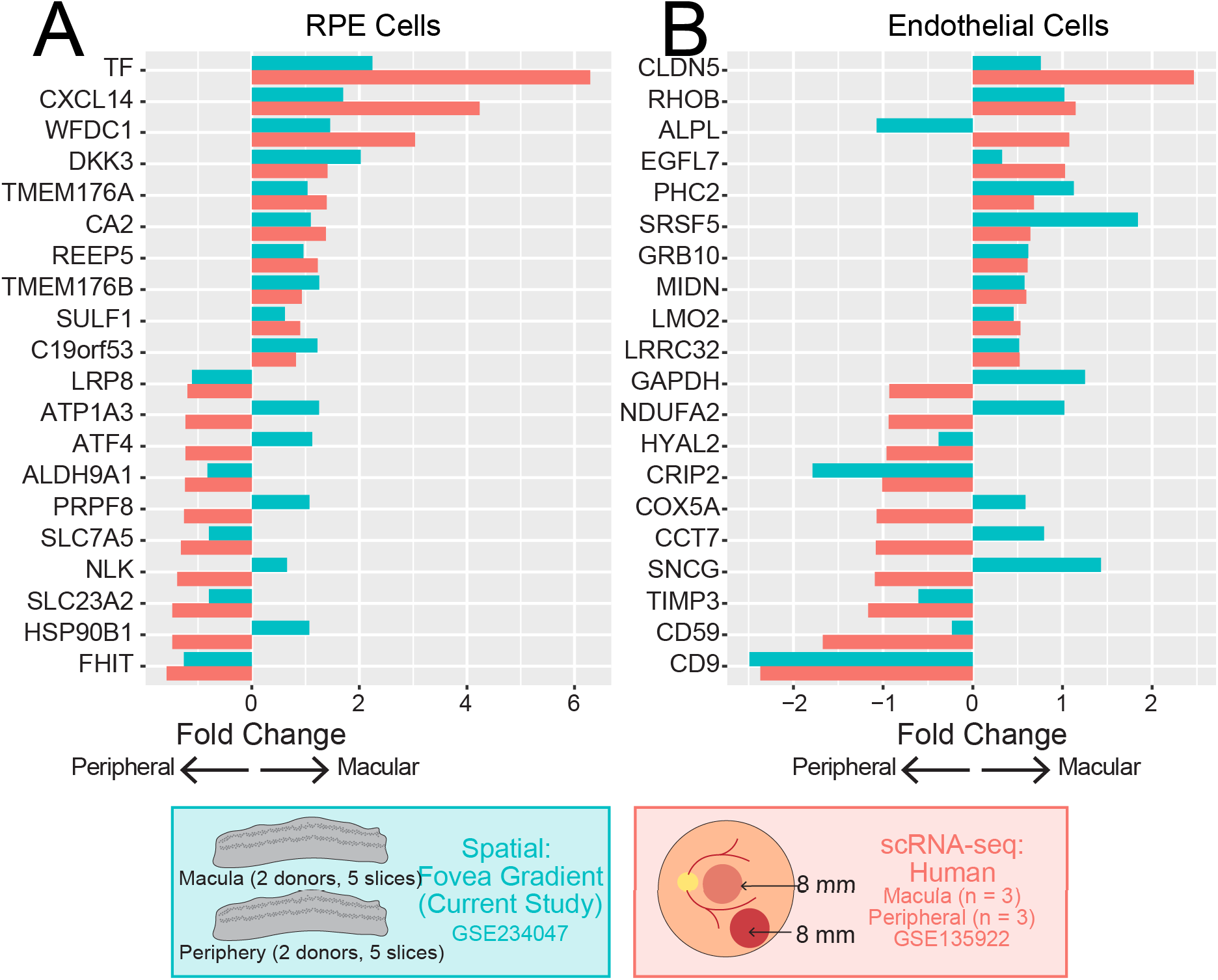
Comparison of macular and peripheral gene expression between spatial and single-cell RNA sequencing studies. Differential expression was completed to compare macular and peripheral enriched genes in the RPE and choroid. Cell type deconvolution and region were used as covariates for differential expression analysis. For RPE cells (**A**) and endothelial cells (**B**), the top differentially expressed genes between the macula and periphery were identified from a previous single-cell RNA sequencing study. Differential expression results were compared between this previous study (red) and the current spatial study (blue). Positive log_2_ fold-changes are associated with increased expression in the macula.

## DISCUSSION

AMD remains a common cause of global blindness. MNV occurs in approximately 5 - 20% of cases (37, 38) and can result in rapid and severe loss of vision. While anti-VEGF therapies have been widely successful at protecting vision in the setting of MNV, resistance to anti-VEGF therapy is common and can reduce the therapeutic effect, leading to visual deterioration (39). Identifying new molecular features of neovascular choroidal endothelial cells may help advance understanding of MNV triggers and provide insight into treatment resistance to guide new potential therapeutic targets.

Bulk (40, 41) and single-cell (13, 14, 36, 42-44) transcriptomic experiments have allowed for the study of gene expression in choroidal endothelial cells across health, AMD, and rodent models of neovascularization. Likewise, single-cell RNA sequencing has been utilized to study gene expression changes in models of retinal degeneration (45, 46). However, MNV, like many other retinal conditions, is a highly regional disease in which focal patterns of pathology are often directly adjacent to more normal retina, RPE, and choroid. It is challenging to use single-cell based approaches to study such heterogenous, spatial disease processes because within each sample, there is no ground-truth classification that labels if cells originate from a healthy versus diseased area. Spatial RNA sequencing addresses this limitation by allowing an expert to label areas of pathology, and this facilitates more specific gene expression comparisons between the lesion and non-diseased areas.

The spatial technology employed in this study had a resolution of 55 micrometers. This spot diameter was not near the single-cell level of resolution, and investigation of H&E images revealed that most spots were covered by several cells and usually represented a mix of different cell types (**Figure 1, 2**). However, this level of resolution still allowed for the investigation of gene expression of different retinal, RPE, and choroidal cell types. For example, known retinal ganglion cell, photoreceptor, and RPE specific genes clearly localized to spots in different retinal layers (**Figure 2C**, **Figure 2E, SI Figure 1**). In addition, a deconvolution algorithm estimated the cell type composition of each spot (**Figure 2**), and retinal layers were easily differentiated by their unique deconvoluted compositions.

Capturing the precise spatial location of each spot also allowed for new differential expression analyses. For example, we identified genes within the neural retina that had varied expression along a continuous gradient from the center of each section (and nearest the fovea) to the edges (**Figure 5**). The identified foveal-enriched genes were highly correlated with previous single-cell level investigations, highlighting the power of spatial technology to perform nuanced biological comparisons. As technology evolves, similar analyses in which the spatial location of gene expression is investigated along a distance gradient may provide insight into the regional predilection of other retinal conditions, such as inherited retinal diseases.

The primary aim of this study was to identify genes dysregulated within the MNV. Although we acquired three different sections through the MNV lesion, with a diameter of 55 microns, no spot was entirely contained within the carefully traced neovascular membrane. As such, applying the CSIDE algorithm allowed us to control for the two major sources of variation in differential expression: cell type composition and MNV overlap. Instead of setting a cutoff value and arbitrarily classifying which spots were in and out of the neovascular membrane, CSIDE permits continuous explanatory variables. Therefore, we used ImageJ to quantify the intersection between each spot and the carefully traced MNV and used this value to identify genes associated with MNV overlap.

MNV-enriched genes expressed by RPE cells and endothelial cells were predicted to belong to known angiogenesis and neovascular pathways (**Figure 3**). For example, the MNV lesion was enriched in integrins (such as endothelial *ITGB1* and RPE *ITGB8*) previously associated with angiogenesis and blood vessel network formation (24, 29). Likewise, VEGF and TGFB1, which have been extensively associated with neovascularization, were identified as two of the most likely upstream regulators of the observed MNV-enriched genes predicted to originate from endothelial cells and RPE cells (**Figure 3 I-J**). In addition, we compared MNV-enriched genes identified by the current spatial study with a previous single-cell experiment of two MNV donors (14) as well as a rodent model of choroidal neovascularization (13). The most upregulated endothelial MNV gene in the current study, *IL6*, has been previously shown to be expressed by choroidal macrophages and associated with neovascularization in rodents (35). Collectively, these findings suggest that at the current resolution, spatial RNA sequencing can identify known pathogenic drivers of MNV and offers a useful approach to study gene expression within this highly spatial disease process.

There are several limitations to this study. First, as mentioned above, the 55-micrometer diameter resolution of the spatial technology captured several cells within each spot. We partially addressed this limitation by using the non-parametric differential expression algorithm CSIDE, which deconvoluted cell type composition within each spot and stratified differential expression results by cell type. However, such deconvolutions are imperfect, and the unambiguous determination of cell types would be improved with greater spatial resolution. Second, only one MNV donor and one age-matched control donor were used in the current study. While this MNV donor represents a valuable sample with correlated clinical imaging, histopathology, and spatial gene expression, additional validation of the observed expression changes in other human donors is required.

Spatial gene expression is a rapidly evolving technology (47). Recently commercialized systems allow for true single-cell or sub-cellular resolution of gene expression within a spatial context (e.g. (48, 49)). However, such technologies use probe-based hybridization and thus necessitate custom gene panel designs. Similarly, unbiased sequencing-based platforms are improving spot resolution to the near single-cell level. This current investigation demonstrates that spatial technology can be used to query gene expression in focal areas of pathology and along regional distance gradients within the heterogenous retina, RPE, and choroid. Further studies, along with technological advances, will continue to improve our understanding of pathogenesis mechanisms in MNV and other regional retinal diseases. Insights from this potent technology may guide the development of future targeted therapeutic interventions with potential to decrease vision loss from AMD and MNV.

---

**SI Figure 1:**
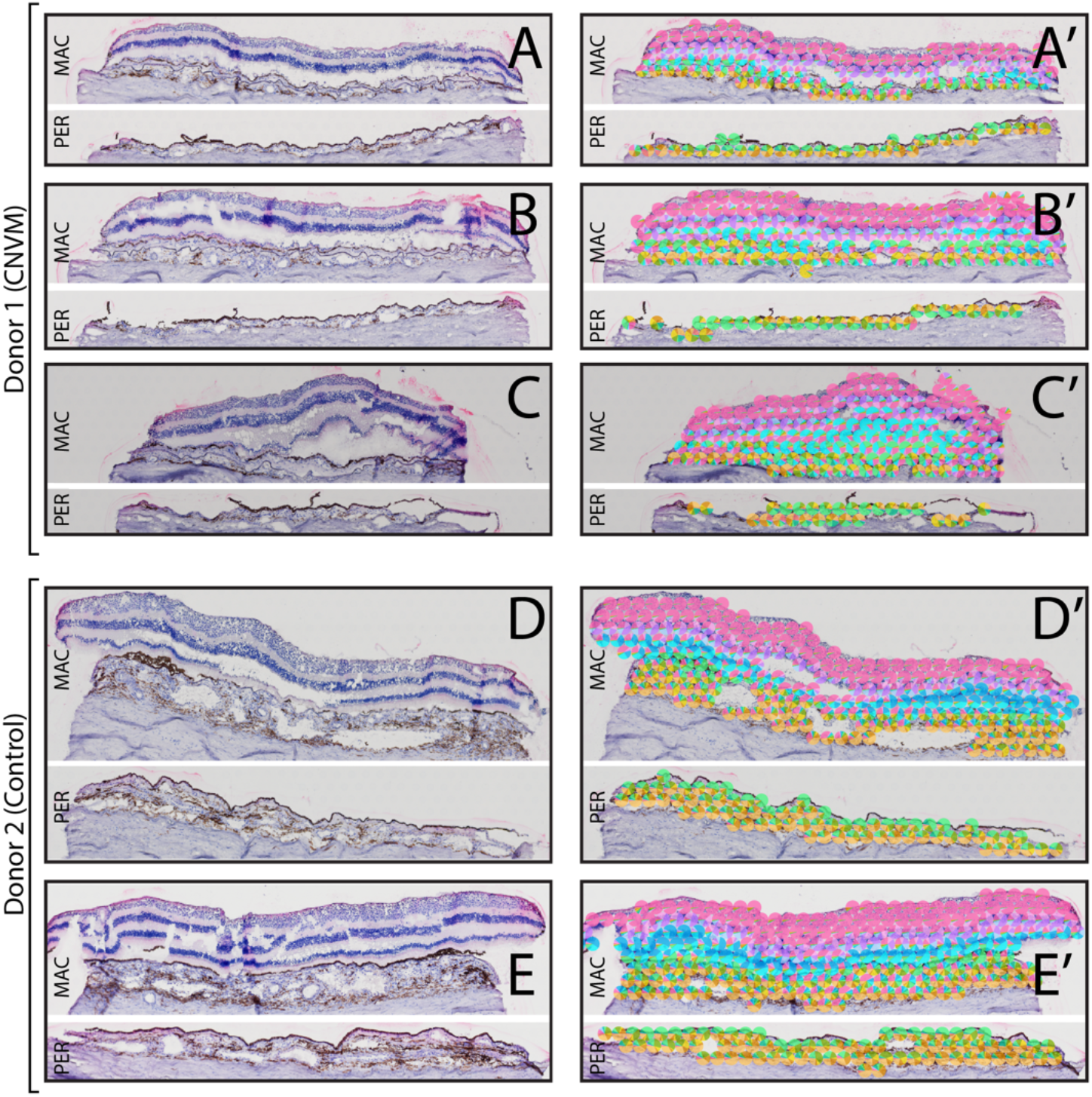
Three H&E tissue sections from the MNV donor (**A-C**) and two tissue sections from the control donor (**D-E**) are visualized. **A’ – E’:** The RNA contribution from retinal, RPE, and choroidal cell types was estimated in each spatially barcoded spot and displayed as a pie-chart as designated in Figure 2.

**SI Figure 2:**
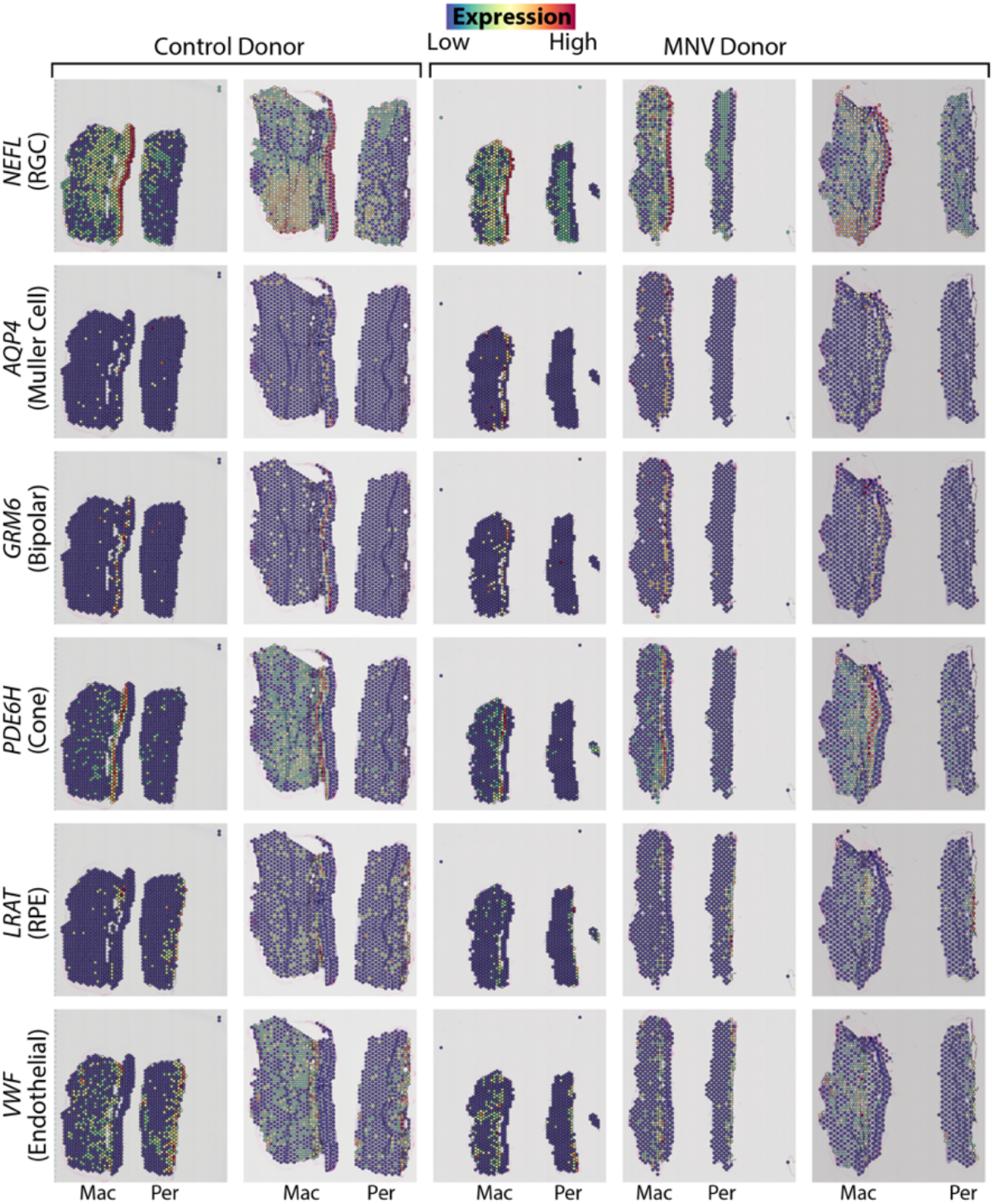
Spatial expression of hallmark retinal, RPE, and choroidal genes. Each row depicts the spatial expression of one gene, with a parenthetical comment of the cell population that most highly expresses that gene (according to previous single-cell RNA sequencing studies). Each column depicts one of the five sections used in the spatial transcriptomics experiments (columns 1-2 correspond to the control donor while columns 3-5 correspond to the MNV donor). Each 55-micron capture spot is outlined in black, and the color of the spot represents the detected expression (red corresponds to high expression while blue corresponds to low expression).

## ACKNOWLEDGEMENTS

The authors gratefully acknowledge the eye donors and their families for their generous contributions to this research, as well as the Iowa Lions Eye Bank for their assistance in providing this donor tissue. Visium data presented herein were obtained at the Iowa NeuroBank Core in the Iowa Neuroscience Institute (INI) and the Genomics Division in the Iowa Institute of Human Genetics (IIHG).

## Notes

### Competing Interest Statement

The authors have declared no competing interest.

